# Precision Editing of Cyclophilin A Generates Cyclosporine and Voclosporin Resistant Cellular Therapies

**DOI:** 10.1101/2025.04.30.651441

**Authors:** Holly Wobma, Francesca Alvarez Calderon, Jiayi Dong, Kayleigh Omdahl, Xianliang Rui, Elisa J. Rojas Palato, Rene S. Bermea, Alexandre Albanese, Franziska Wachter, Marlana Winschel, Katherine A. Michaelis, William A. Burns, Gillian Selig, Lorenzo Cagnin, Victor Tkachev, Susan E. Prockop, Peter A. Nigrovic, Bruce R. Blazar, Ulrike Gerdemann, Leslie S. Kean

## Abstract

Recipients of allogeneic transplants or patients with autoimmune disease require immune suppression, often with calcineurin inhibitors. There is an expanding repertoire of immune effector cell therapies, including CD19 CAR-T cells and viral-specific T cells, deployed in these patients; however, ongoing calcineurin inhibition may be detrimental to cell therapy function. We developed a CRISPR/Cas9-based approach to engineer dual cyclosporine/voclosporin resistant cell therapies by targeting *PPIA* (encoding cyclophilin A), a critical binding partner for both drugs. Because Cyclophilin A has homeostatic functions in T cells, a complete knock-out is detrimental to cell viability. We thus targeted its C-terminus, disrupting drug binding while leaving the majority of the protein intact. C-terminal editing was stable throughout expansion and preserved Cyclophilin A expression. Edited CD19 CAR-T cells retained effector function in the presence of cyclosporine and voclosporin, including proliferation, cytokine production, and target cell killing, resulting in improved survival in murine models of CD19+ leukemia. Edited CMV-specific T cells also demonstrated preserved antigen-specific proliferation and cytokine production in the presence of these drugs. C-terminal editing of Cyclophilin A offers a promising avenue for developing next-generation cell therapies for patients receiving calcineurin inhibitors.

## Introduction

Immunosuppression is the cornerstone of management of allo-and auto-immunity. Calcineurin inhibitors (CNIs) are a key medication class used to prevent graft-vs-host-disease after hematopoietic cell transplantation (HCT),^1^ to prevent graft rejection after solid organ transplantation (SOT),^2^ and to treat autoimmune disease.^3^ Standard CNIs include cyclosporine (CsA) and tacrolimus, both widely used in HCT and SOT. Next-generation CNIs with more favorable pharmacokinetic properties are also being developed, including the CsA derivative, voclosporin (VCS), which is FDA approved for patients with lupus nephritis, expanding application of this medication class.^4^

A key attribute of CNIs is their ability to inhibit lymphocyte proliferation, persistence, and function. While these characteristics form the basis of their efficacy, they can be disadvantageous for patients with allo-or auto-immune disease experiencing complications such as relapse and infection, who are increasingly being treated with adoptive immune effector cells (IEC). These IECs include chimeric antigen receptor (CAR)-T cells and viral-specific T cells (VSTs), each of which may be inhibited by CNIs. Managing the delivery of IECs in these patients is complex, as CNI withdrawal risks exacerbating allo-or auto-immunity.^5^ Engineering CNI-resistance is one way to optimize the potential of IEC therapies.

Calcineurin is a phosphatase that becomes active with binding of calcium/calmodulin following T cell receptor (TCR) activation. Calcineurin mediates T cell activity in two ways: 1) it dephosphorylates the transcription factor NFAT, which can then enter the nucleus and induce expression of genes related to T cell activation (pro-inflammatory cytokines, co-stimulatory proteins),^6^ and 2) it removes an inhibitory phosphate from LCK that helps maintain upstream TCR signaling. Both mechanisms further contribute to immune synapse formation and enhanced cytotoxicity.^7^ Additionally, calcineurin dephosphorylates synaptopodin in the kidney, resulting in podocyte instability and increased proteinuria in lupus.^8^ Thus, the therapeutic benefit of CNIs derives from inhibition of T cell activation, with disease-and organ-specific effects also observed.

CNIs rely on immunophilins to bind calcineurin and exert their inhibitory effect.

Immunophilins are a family of peptidyl-prolyl isomerases that are highly evolutionarily conserved and widely expressed.^9^ They can be divided into ‘FK506 binding proteins’ (FKBP) and cyclophilins. Since tacrolimus and CsA/VCS can bind to calcineurin only after pairing with FKBP12 and Cyclophilin A (CypA), respectively,^10^ eliminating their expression represents a rational approach for engineering CNI resistance. Indeed, tacrolimus resistance has been achieved by knocking-out FKBP12.^11–15^ In this case, knock-out also impacts the mTOR pathway, conferring resistance to rapamycin that is inseparable from resistance to tacrolimus. Unlike for FKBP12, knock-out of CypA in T cells has not been reported. This is likely due to the widespread homeostatic functions of CypA in immune cells such that complete deletion may be incompatible with T cell survival and function.^16–18^

To engineer CsA/VCS-resistant IECs, we therefore employed selective CRISPR/Cas9 editing to induce a frameshift in the last exon of *PPIA* (encoding CypA), thereby disrupting the CNI interaction site, and thus the inhibitory effect of CNIs on calcineurin, while leaving the core CypA structure intact. Here, we demonstrate this targeted approach leads to resistance to both CsA and VCS in otherwise fully functional CAR-T cells and VSTs.

## Results

### C-Terminal PPIA editing retains CypA expression

To confer CNI resistance to T cells, we selectively disrupted the C-terminal region of CypA, which is critical for the interaction of CsA/VCS-CypA with calcineurin A.^19^ This was achieved via targeted CRISPR/Cas9 editing of the last exon (Exon 5) of *PPIA* in T cells from healthy donors (**Fig. 1A**), a strategy that maintained CypA protein expression and T cell viability. As a comparator, we edited the immediate upstream exon (Exon 4, **Fig. 1A**), an approach predicted to lead to complete gene knock-out (NMDetective-A^20^ score 0.78, with scores >0.52 corresponding to translation products predicted to undergo non-sense mediated decay (NMD)).^20^ With the Exon 4 edit, we observed high levels of early cell death compared to controls (**Fig. 1B)**, rapid loss of the edited T cells during cell culture **(Fig. 1C),** and consequently, minimal proliferation of edited cells **(Fig. 1D).** Notably, the Exon 5 approach was associated with a low NMDetective-A score (0.23), low early cell death, and comparable growth kinetics to *PPIA^WT^* cells **(Fig. 1B-D),** including an average of 3.9 population doublings in the first 10 days after CRISPR editing, supporting the feasibility of this engineering approach. Given the promising results with our Exon 5 edit, and the predicted C-terminal disruption, we henceforth refer to cells with the Exon 5 edit as ‘*PPIA*^Δ*C*^*’*.

**Figure 1.**
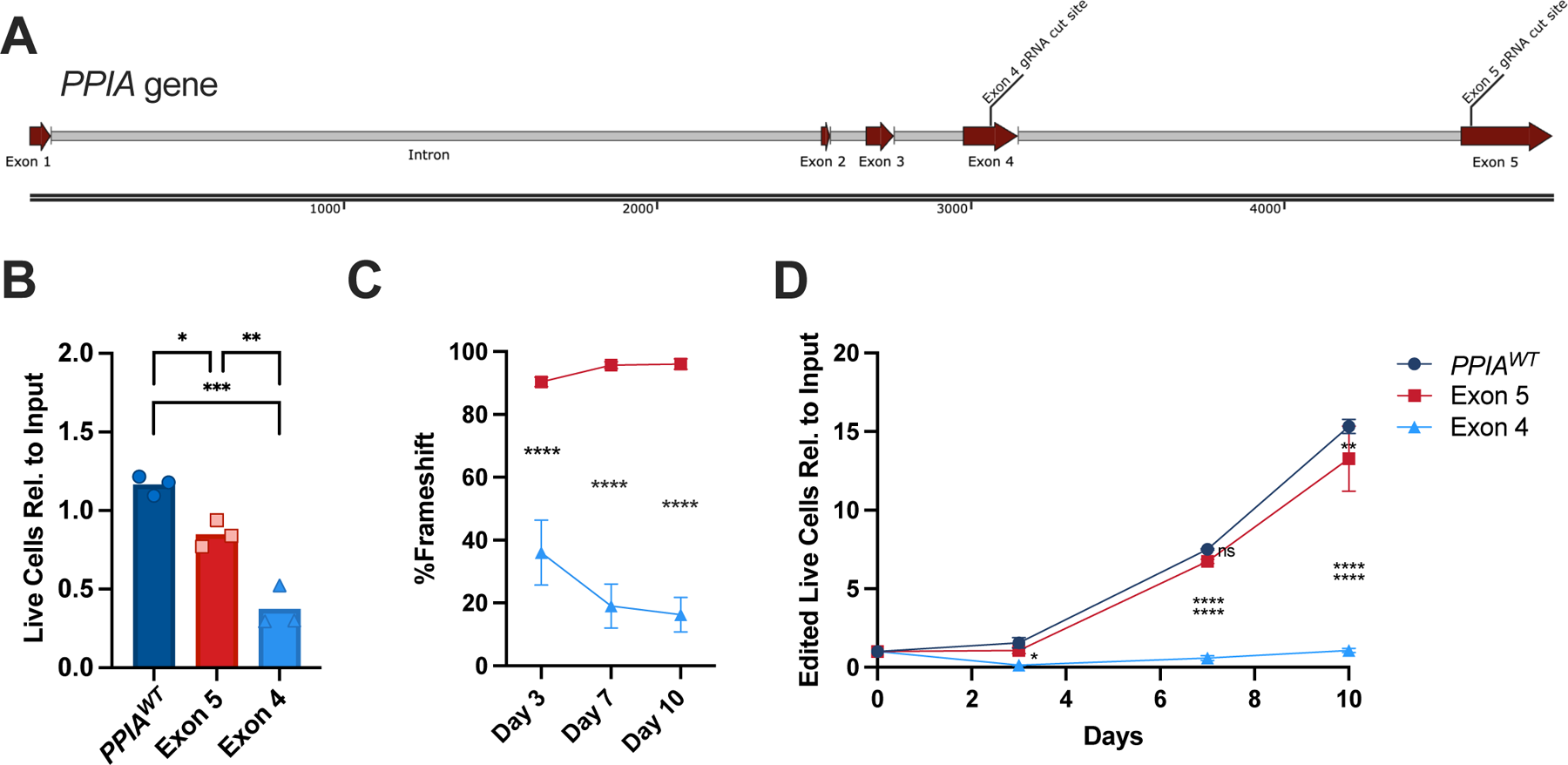
CRISPR/Cas9 editing of Exon 5 but not Exon 4 facilitates expansion of edited cells. A) Schematic of the *PPIA* gene containing 5 exons, the largest of which are Exon 4 and Exon 5. CRISPR/Cas9 gRNA cut sites are indicated, and guide sequences are provided in the Methods. B) Significantly lower live T cell counts relative to the number of electroporated cells (input) after CRISPR/Cas9 targeting of Exon 4, two days after gene editing. Control, *PPIA^WT^*cells were edited with a non-targeting guide. Groups compared via One-way ANOVA with Tukey’s post-hoc test. C) Change of Exon 4 and Exon 5 editing rates over time (relative to date of CRISPR/Cas9 RNP nucleofection) as determined by Sanger sequencing on Days 3 (first time point of assessing editing rate), 7 and 10. D) The degree of expansion of CRISPR-Cas9 edited cells was calculated based on the live cell count multiplied by % edited cells relative to the Day 0 input cell count. The * at Day 3 reflects the comparison between *PPIA^WT^*and the Exon 4 edit. The remaining Day 3 comparisons were not significantly different (ns). Experiments performed with n = 3 donors per group; data represent mean (SD); error bars are not visible when smaller than the marker size. *<0.05, **<0.01, ***<0.001, ****<0.0001

*PPIA*^Δ*C*^ cells demonstrated a high editing efficiency 89.5 (1.3)% measured on day 3 (**Fig. 2A-B**), predominantly comprised of a single nucleotide insertion (+1 base pair (bp) frameshift) in 58 (12)% of cells, followed by a single nucleotide deletion (-1 bp frameshift) in 26 (8%) of cells (**Fig. 2C**). This editing efficiency and indel distribution was highly consistent across donors, and edited cells demonstrated a stable editing efficiency and indel distribution over 3-4 weeks in culture (**Fig. 2C**), indicating compatibility with cell survival. As hypothesized, CypA protein expression was preserved when assessed at day 10, as demonstrated by flow cytometry, Western blot, and confocal imaging (**Fig. 2D-F).** To confirm long term CypA protein expression after *PPIA*^Δ*C*^ editing, Jurkat cells containing the dominant +1 bp indel were expanded for >8 weeks in culture demonstrated preserved editing efficiency (98% +1 bp indel) and CypA protein expression (**Suppl. Fig. 1A**).

**Figure 2.**
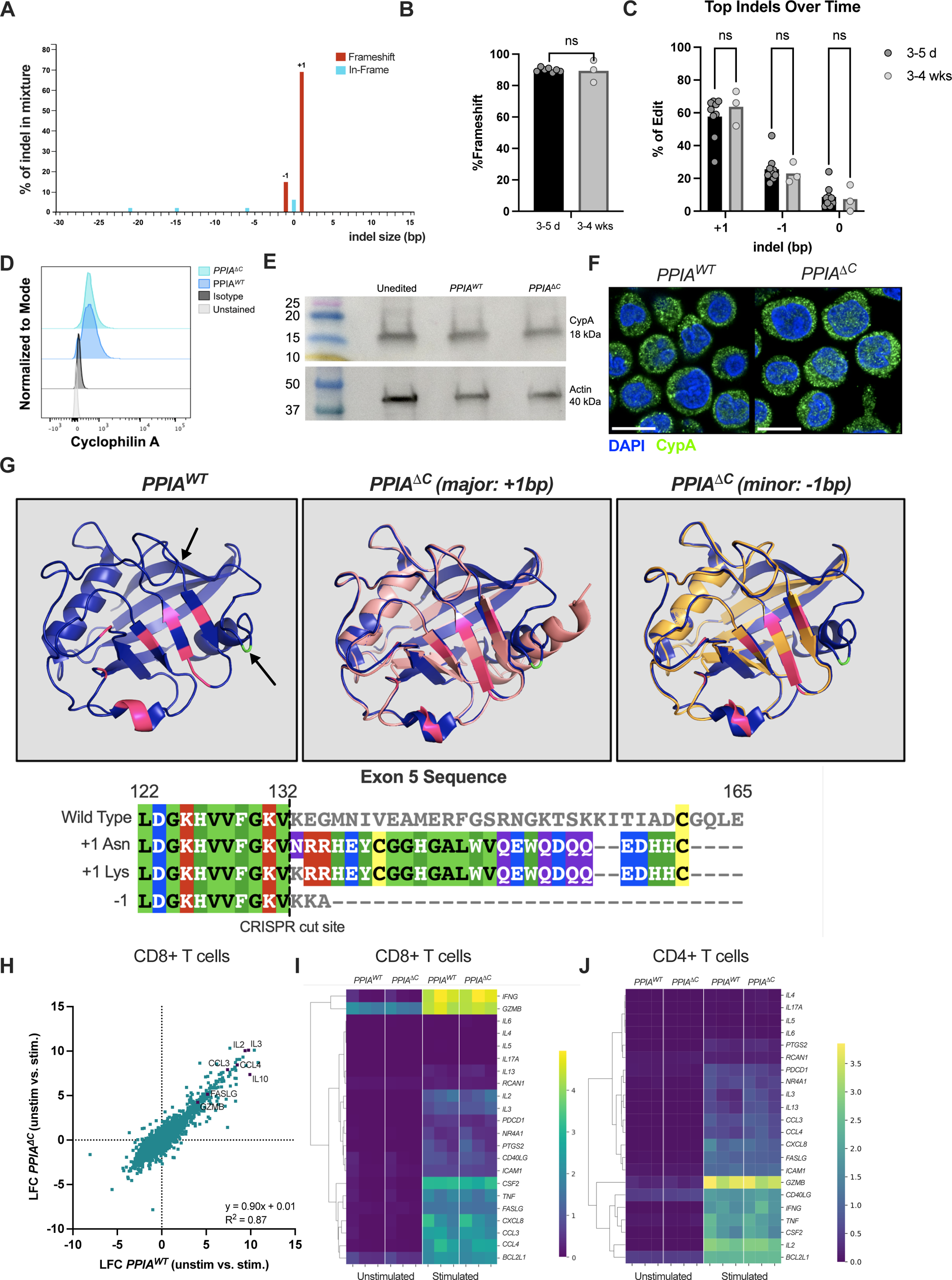
Characterizing the *PPIA*^Δ*C*^ edit and associated protein. A) CRISPR editing at a specific locus in Exon 5 leads to a consistent indel distribution of mostly +1 >-1 bp indels. B-C) The overall editing efficiency and indel distribution are stable over 3-4 weeks (n = 3-9). D-F) CypA protein expression 10 days after CRISPR/Cas9 editing by (D) flow cytometry and (E) Western blot using actin as a loading control (after stripping the membrane and re-staining) and a Precision Plus Protein^TM^ Kaleidoscope^TM^ ladder with corresponding protein sizes indicated on the left; no image enhancement software was used. (F) Confocal imaging also confirming intracellular presence of CypA. Scale bar represents 10 µm. G) Predicted 3D protein structure based on AlphaFold Colab where the edited protein (rose or tangerine) is overlayed on the wildtype protein (purple). Catalytic residues important to homeostatic function are highlighted in pink (R55, F60, M61, Q63, A101, F113, W121, L122, and H126). An important residue for CsA binding is shown in green (R148; see black arrow). Another black arrow points towards a f3-strand absent in the edited proteins. The corresponding amino acid sequences in Exon 5 are shown underneath as well as the CRISPR cut site after amino acid 132. H) Comparison of the log fold change (LFC) in expression (unstimulated vs. stimulated) in *PPIA^WT^* and *PPIA*^Δ*C*^ CD8+ T cells, analyzed by a simple linear regression model. p<0.0001. Several NFAT-inducible genes are shown with purple markers. H-I) Normalized gene expression based on bulk RNA-seq in *PPIA^WT^* and *PPIA*^Δ*C*^ (I) CD8+ T cells and (J) CD4+ T cells, highlighting genes related to TCR activation and the NFAT signaling pathway. n = 3 donors. *<0.05, **<0.01, ***<0.001, ****<0.0001

Based on the two predominant indels, we modeled predicted *PPIA*^Δ*C*^ protein structures using AlphaFold Colab^22,23^ and aligned them with the wildtype protein (*PPIA^WT^* shown in purple; **Fig. 2G**). As expected, the two predicted *PPIA*^Δ*C*^ protein structures align around important residues for CypA function (pink)^19,24^ but vary near Arg148 (green),^19^ an amino acid critical for the interaction of CypA with the CsA-calcineurin complex.^19^ For the +1 bp indel protein, the regional loss of structural homology was due to a frameshift after amino acid 132, with the protein only minimally truncated (159 amino acids vs. wildtype of 165). For the-1 bp indel, the frameshift led to a premature stop codon at amino acid 135 and thus a protein truncated prior to Arg148.

To verify that cells with *PPIA*^Δ*C*^ retained wildtype T cell transcription patterns as well as functional TCR signaling pathways – critical for any IEC therapy – we performed bulk RNA-seq on purified CD8+ and CD4+ T cells expressing either *PPIA^WT^* (mock edited) or *PPIA*^Δ*C*^ at baseline and upon stimulation with anti-CD3/CD28 Dynabeads. Unlike the significant transcriptional differences between edited and wildtype cells previously reported for gene-addition approaches to CsA resistance^25^, the transcriptomes of the *PPIA^WT^* and *PPIA*^Δ*C*^ CD8+ T cell populations were highly similar, with no differentially expressed genes, either at baseline or after stimulation **(Fig. 2H-I, Suppl. Fig. 1B)**. This included comparable induction of NFAT pathway related genes (CCL3, CCL4, IL2, TNF-a, IFN-y) upon stimulation in both *PPIA^WT^* and *PPIA*^Δ*C*^ cells (**Fig. 2H-I**). Similar bulk RNA-seq results were obtained from CD4+ T cells (**Fig. 2J**). Finally, *PPIA*^Δ*C*^ cells adopted similar maturation patterns (naïve, central memory, effector memory) to *PPIA^WT^* cells after bead stimulation (**Suppl. Fig. 1C**).

### PPIA^ΔC^ T cells are resistant to CsA and VCS

To determine whether *PPIA*^Δ*C*^ cells were resistant to CsA and VCS, we performed a mixed lymphocyte reaction comparing division of *PPIA^WT^* and *PPIA*^Δ*C*^ responder T cells in the presence of increasing doses of each agent (**Fig. 3**). This dose-escalation importantly revealed that lower CsA doses are inhibitory *in vitro* relative to those commonly targeted in patients. As shown in **Fig. 3C**, *PPIA^WT^* responder T cells exhibited a dose-dependent suppression of proliferation, while *PPIA*^Δ*C*^ responder T cells were substantially resistant to CsA/VCS suppression across all doses. For example, at a dose of 25 ng/mL CsA, *PPIA^WT^* responder CD4+ T cells demonstrated 69 (8)% suppression of division, whereas *PPIA*^Δ*C*^ responder CD4+ T cells demonstrated less suppression (28 (12)%, p < 0.0001). Similarly, at a dose of 10 ng/mL VCS, *PPIA^WT^* responder CD4+ T cells demonstrated 84 (5)% suppression of division by drug, whereas *PPIA*^Δ*C*^ responder CD4+ T cells demonstrated less suppression (42 (8)%, p < 0.0001). These results were similar for CD8+ responder T cells. In contrast, and consistent with the dependency of tacrolimus and rapamycin on the alternative immunophilin, FKBP12, division of *PPIA*^Δ*C*^ and *PPIA^WT^* responder T cells were equally suppressible by tacrolimus and rapamycin (**Fig. 3D**).

**Figure 3.**
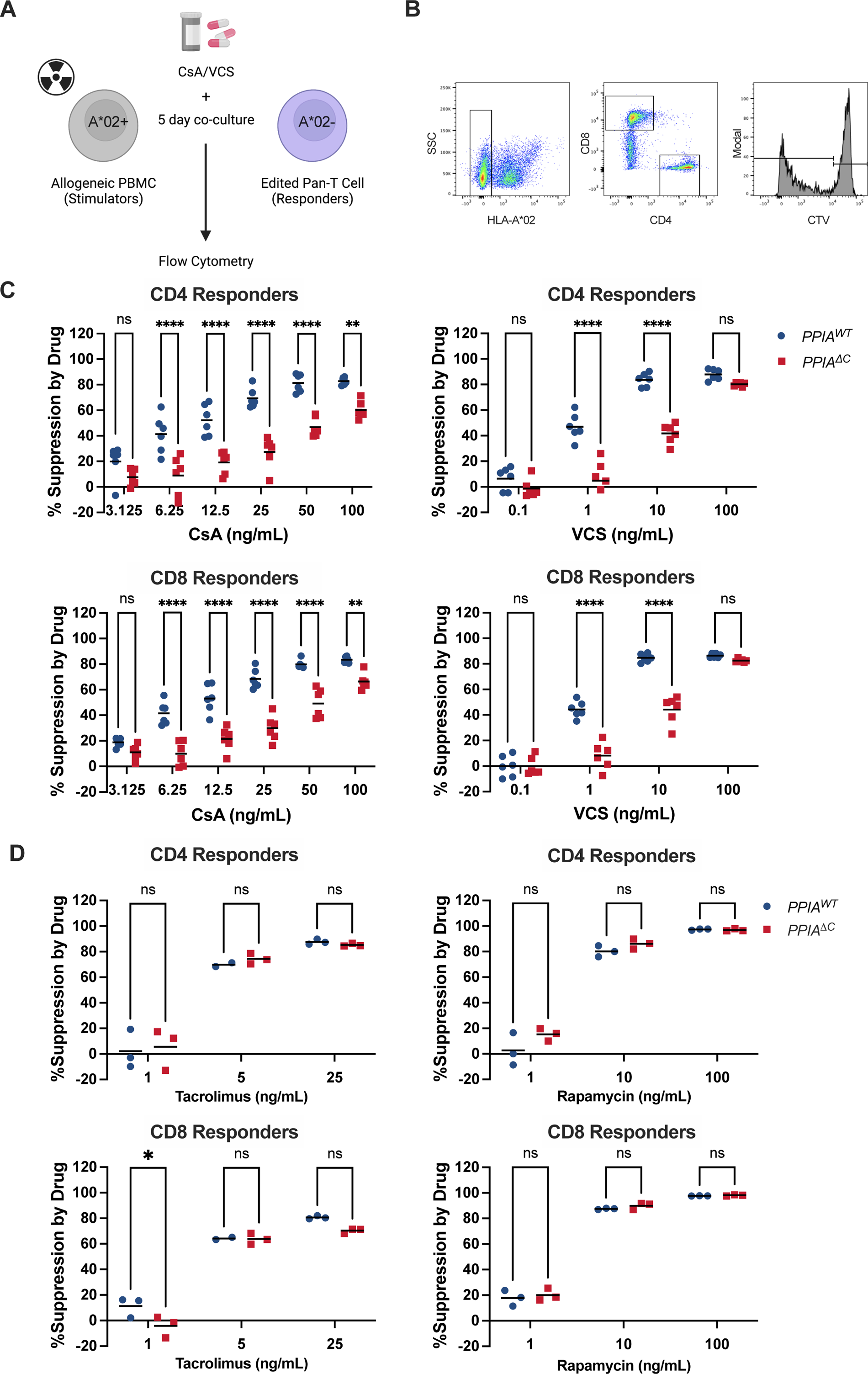
*PPIA*^Δ*C*^ T cell division is resistant to CsA and VCS suppression. A) Schematic of a mixed lymphocyte reaction suppression assay in which ‘stimulator’ PBMCs are in co-culture with ‘responder’ *PPIA^WT^* or *PPIA*^Δ*C*^ T cells at a 3:1 ratio B) Gating strategy of assessing proliferating cells in the mixed lymphocyte reaction: Stimulator and responder cells were HLA-A*02 disparate allowing separation of populations on flow cytometry. Responder cells were gated on CD4+ and CD8+ T cell subsets and proliferation was evaluated based on CTV dilution within these populations. C-D) Division of Responder CD4+ and CD8+ T cells was assessed based on CTV dilution relative to a “Responder only” negative control group across a (C) CsA and VCS dose escalation and (D) tacrolimus and rapamycin dose escalation. Percent suppression by drug was calculated by normalizing data to the drug-free (positive) control (see Methods). Black bars show the mean. 2-way ANOVA with Sidak post-hoc test was used for statistical analysis. n = 6 replicates. *<0.05, **<0.01, ***<0.001, ****<0.0001

### PPIA^ΔC^ CD19 CAR-T cell activation and proliferation is resistant to CsA and VCS

Having established CsA and VCS resistance in primary T cells, we next sought to validate drug resistance in representative IEC therapies, beginning with CD19 CAR-T cells. Current FDA-approved CD19 CAR-T cell products contain either 41BB or CD28 costimulatory domains (denoted BB(and 28(, respectively).^26^ Given the distinct signaling pathway these costimulatory molecules confer, we first evaluated the ability of CsA to inhibit proliferation of BB(and 28(CAR-T cells in *PPIA^WT^* CAR-T using CD19+ NALM6 f3*2M^KO^* cells as stimulators. As shown in **Fig. 4**, both BB(and 28(were suppressible by CsA but exhibited different drug sensitivities, with BB(-CAR-T more sensitive to CsA inhibition than 28(-CAR-T cells. For example, at a low effector-to-target (E:T) ratio (1:1), there was 6 (1)% suppression of BB(CD8+ CAR-T cell division at the lowest CsA dose (12.5 ng/mL) and 74 (5)% suppression at the highest dose (100 ng/mL). For 28(CD8+ CAR-T cells at the 1:1 E:T ratio, there was no suppression at either drug dose (6 (1)% and-6 (6)% respectively). However, high E:T ratios (10:1) rendered 28(CAR-T cells inhibitable by CsA, with 46 (7)% suppression at a CsA dose of 25 ng/mL and 81 (9)% suppression at a dose of 50 ng/mL. Trends were similar when CD4+ CAR-T cells were assessed (**Suppl. Fig. 2**).

**Figure 4.**
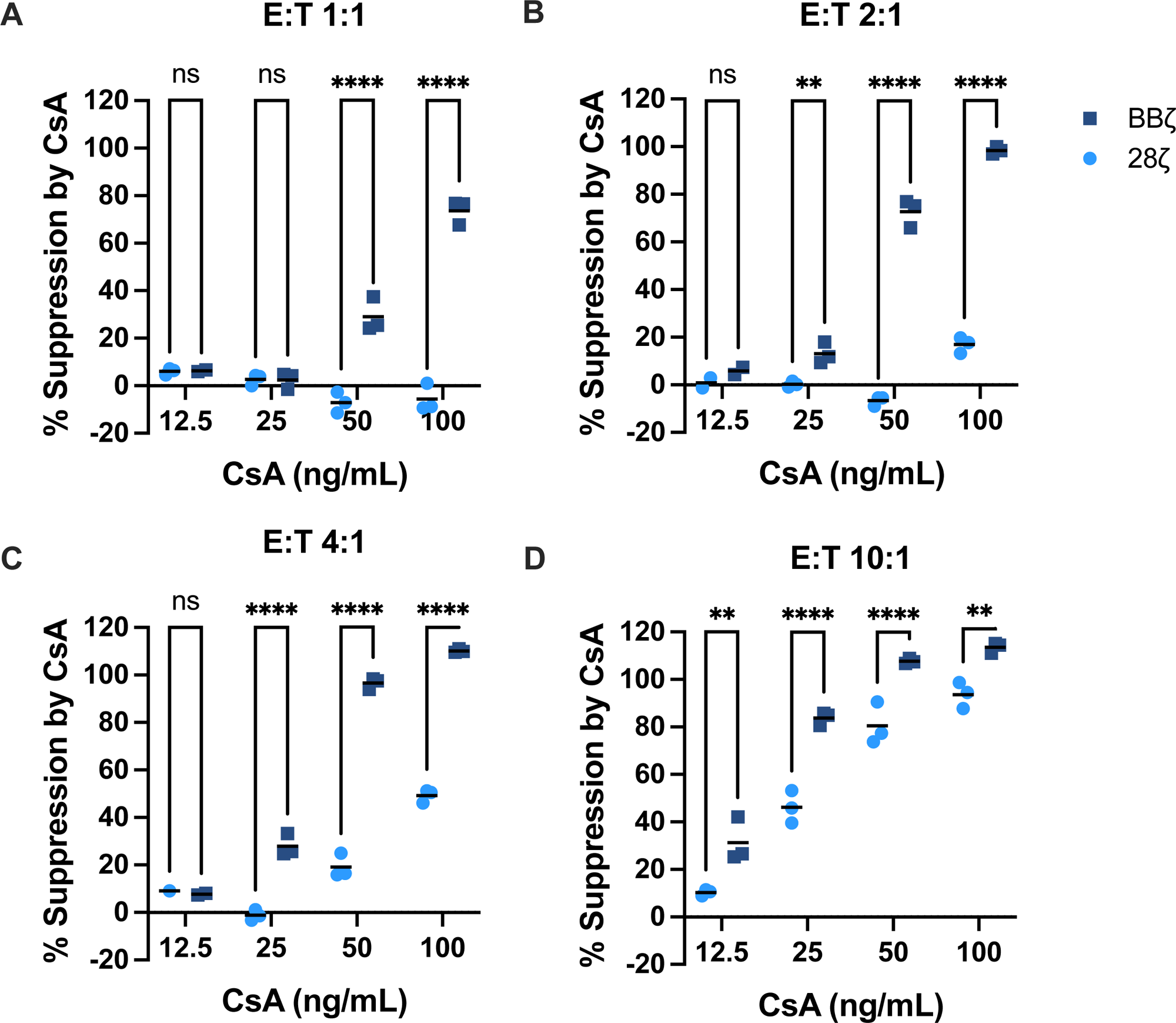
BB**{** CAR-T cells are more sensitive to inhibition by CsA than 28**{** CAR-T cells. Division of CAR-T cells in response to CD19+ f32M^KO^ NALM6 stimulator cells was assessed based on CTV dilution on Day 3, relative to unstimulated CAR-T cells (negative control). The percent of divided cells was normalized to the drug-free (positive) control after subtracting non-specific proliferation in the CAR-only (no tumor) negative controls for each construct (see Methods. Panels are shown for different E:T ratios of A) 1:1 B) 2:1 C) 4:1 and D) 10:1 with a CsA dose titration. Data shown reflects CAR-T cells gated on CD8. Black bars show the mean. 2-way ANOVA with Sidak post-hoc test used for statistical analysis. *<0.05, **<0.01, ***<0.001, ****<0.0001

Next, we evaluated the proliferative potential of CRISPR/Cas9 *PPIA*^Δ*C*^ edited 28(CAR-T cells in response to stimulation with NALM6 *B2M^KO^* cells (E:T 10:1). In vitro the *PPIA*^Δ*C*^ edit in CAR-T cells led to near complete resistance to inhibition of proliferation by CsA and VCS (**Fig. 5A-B** and **Suppl. Fig. 3).** For example, at a CsA dose of 50 ng/mL, there was 45 (8)% suppression of division of CD4+ *PPIA^WT^* CAR-T cells vs. 3 (1)% suppression of *PPIA*^Δ*C*^ cells (p < 0.0001). Similar findings were shown across different doses when cells were co-cultured in the presence of VCS – for example, at a dose of 10 ng/mL, there was 41 (5)% suppression of division of CD4+ *PPIA^WT^* CAR-T cells vs. 1 (2)% suppression of *PPIA*^Δ*C*^ cells (p < 0.0001).

**Figure 5.**
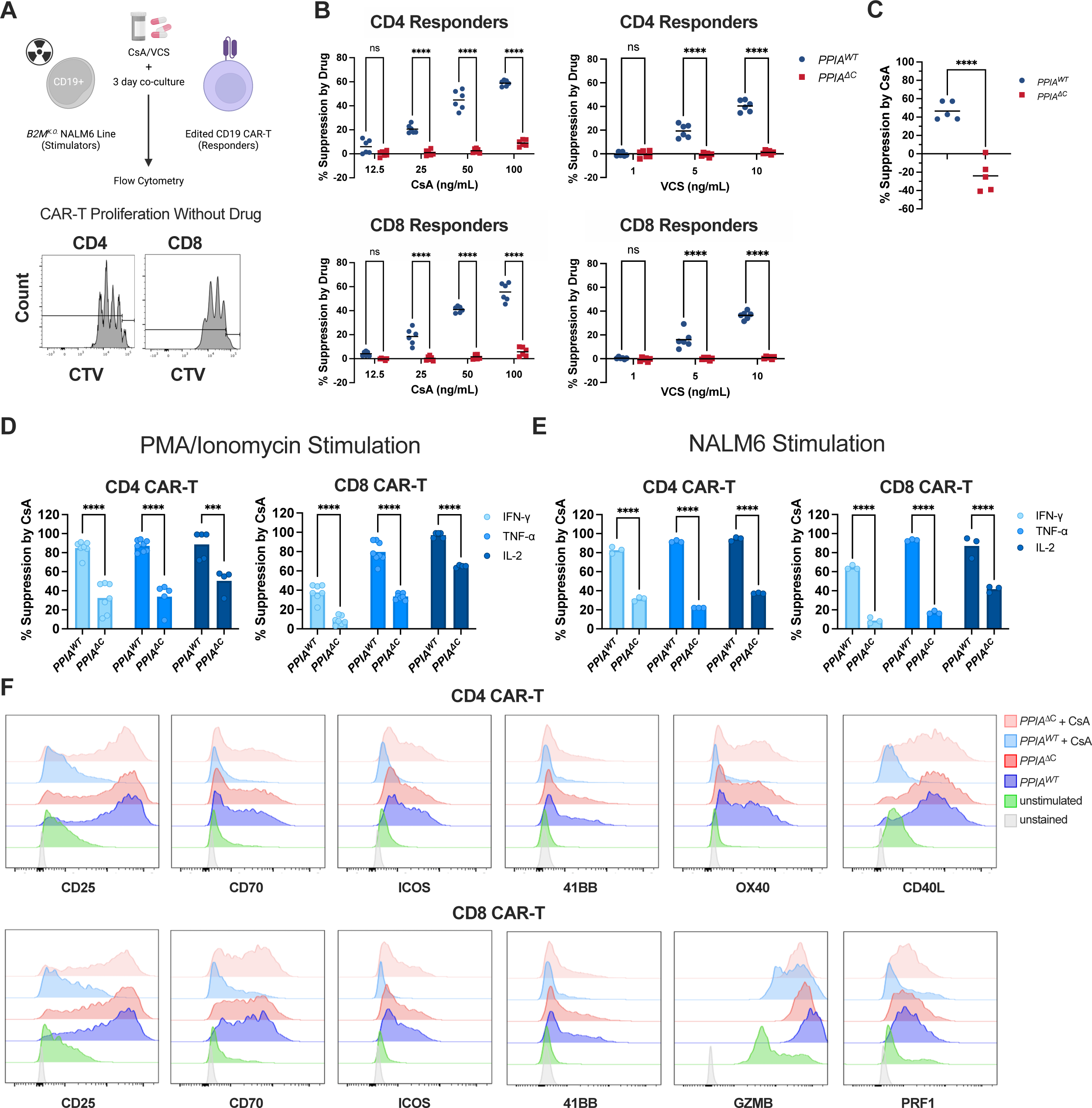
*PPIA*^Δ*C*^ CAR-T cells exhibit preserved division, cytokine production, induction of NFAT-inducible genes in the presence of drug. A) Edited CD19 CAR-T cells were stimulated by f32M^KO^ NALM6 cells at a 10:1 E:T ratio. Division of CAR-T cells was assessed based on CTV dilution on Day 3, comparing to unstimulated CAR-T cells (negative control). The percent of divided cells was normalized to the drug-free (positive) control (see Methods). B) Data shown for CD4+ and CD8+ CAR-T cell division across a CsA and VCS dose titration. Black bars show the mean. n = 6 replicates. 2-way ANOVA with Sidak post-hoc test used for statistical analysis. C) CTV labelled CD19 CAR-T cells were injected into immune deficient (NSG) mice previously inoculated with human CD19+ NALM6 cells. Mice received daily subcutaneous injections of CsA or PBS. Four days after infusion, CAR-T cell division was assessed based on CTV dilution after gating on murine CD45-, human CD45+ cells. A two-tailed T-test was used for statistical comparison (n = 5 per group). D-E) To assess cytokine production, *PPIA^WT^* or *PPIA*^Δ*C*^ CAR-T cells were stimulated for 4 hours in the presence of protein transport inhibitor and either (D) PMA/ionomycin or (E) CD19+ NALM6 cells (E:T of 2:1), and stained for intracellular IFN-y, TNF-a, and IL-2. 100 ng/mL CsA was used for experiments. The % Suppression was calculated as described in Methods. F) CD19 CAR-T cells were co-cultured with NALM6 cells at a 4:1 ratio for 2 days and either surface or intracellularly stained for the indicated proteins in addition to CD4 or CD8 for initial gating.

Similar results were found for CD8+ CAR-T cells. To assess if *PPIA*^Δ*C*^ 28(CAR-T cells could overcome inhibition of proliferation by CsA *in vivo*, NSG mice were injected with NALM6 cells to provide an antigenic stimulus (**Fig. 5C**). As observed *in vitro*, there was marked inhibition of division of *PPIA^WT^* CAR-T cells in mice treated with CsA, relative to mice treated with PBS (47 ± 10% inhibition), whereas there was no inhibition of division by CsA for the *PPIA*^Δ*C*^ CAR-T cells (-24 ± 17% inhibition, with the negative value indicating preserved expansion during the experiment, p < 0.0001).

To further define the impact of drug and drug resistance on CAR-T cell activation, we evaluated cytokine production in response to a mitogen (PMA/ionomycin) or exposure to the CAR target (NALM6 cells). Without addition of CsA, the fraction of cytokine-producing cells was similar between *PPIA*^Δ*C*^ and *PPIA^WT^* CAR-T cells (**Suppl. Fig. 4**); however, in the presence of 100 ng/ml CsA, *PPIA*^Δ*C*^ CAR-T cells demonstrated sustained production of IFN-y, TNF-a, and IL-2 when compared with *PPIA^WT^* cells in both the CD4+ and CD8+ fractions. For example, in the setting of mitogen stimulation (**Fig. 5D**), there was 38 (8)% suppression of IFN-y induction in *PPIA^WT^* CD8+ T cells in the presence of CsA, and less suppression of IFN-y in the presence of CsA for the *PPIA*^Δ*C*^ CAR-T cells (9 (4)% p < 0.0001). Similar results were observed for TNF-a, with 80 (10)% suppression of TNF-α induction in *PPIA^WT^* CD8+ CAR-T cells versus 33 (3)% (p < 0.0001) for the *PPIA*^Δ*C*^ CAR-T cells. For IL-2, there was 89% (10%) suppression of IL-2 induction in *PPIA^WT^* CD4+ CAR-T cells, versus 50 (13)% (p < 0.0001) for the *PPIA*^Δ*C*^ CAR-T cells. These patterns were recapitulated when edited CAR-T cells were co-cultured with NALM6 cells as a source of stimulation more specifically through the CAR (**Fig. 5E**).

While CsA inhibition of the NFAT pathway is commonly thought to mediate immune suppression through disrupting cytokine production as above, there are many additional NFAT-inducible genes that contribute to T cell activation related to costimulation and effector function. We thus sought to determine the impact of CsA on CAR-T cell activation more broadly. As seen in **Fig. 5F**, after two day co-culture with NALM6 cells (4:1 E:T), both CD4+ and CD8+ CD19 CAR-T cells induced upregulation of CD25, CD70, and ICOS, while CD4+ CAR-T cells also induced CD40L and OX40, and CD8+ CAR-T cells increased expression of granzyme B (GZMB) and perforin (PRF1). This induction was uniformly suppressed in the presence of 100 ng/mL CsA in *PPIA^WT^* CAR-T cells but minimally – or not at all – in *PPIA*^Δ*C*^ CAR-T cells.

### PPIA^ΔC^ CD19 CAR-T cells have superior tumor killing in the setting of CsA

We next evaluated CD19 CAR-T cell cytotoxicity towards two CD19 high-expressing leukemia/lymphoma cell lines – wildtype NALM6 cells (NALM6 WT) and Raji cells (**Suppl. Fig. 5A**). In a 3-day cytotoxicity assay, there was CsA-mediated suppression of CAR-T cell cytotoxicity towards both cell lines (**Fig. 6A**). To assess if *PPIA*^Δ*C*^ CAR-T cells were also able to kill leukemia cells with low CD19 antigen density, we assessed day 3 %killing of NALM6 cells with low CD19 expression (NALM6 CD19^lo^ (**Fig. 6A**)). In fact, in this experiment, *PPIA*^Δ*C*^ CAR-T cells overcame CsA-induced suppression even by the 24-hour mark, and this enhanced cytotoxicity persisted through day 3, as reflected by lower live NALM6 CD19^lo^ tumor burden over time (**Fig. 6B**). The long-term cytotoxic advantage of *PPIA*^Δ*C*^ CAR-T cells in the presence of CsA was also observed when they were cocultured with NALM6 WT cells for up to 7 days, with tumor rechallenge half-way through (**Suppl Fig. 5B**). This sustained cytotoxic effect reflects a combination of preserved effector function and proliferation of *PPIA*^Δ*C*^ CAR-T cells in the presence of CsA.

**Figure 6.**
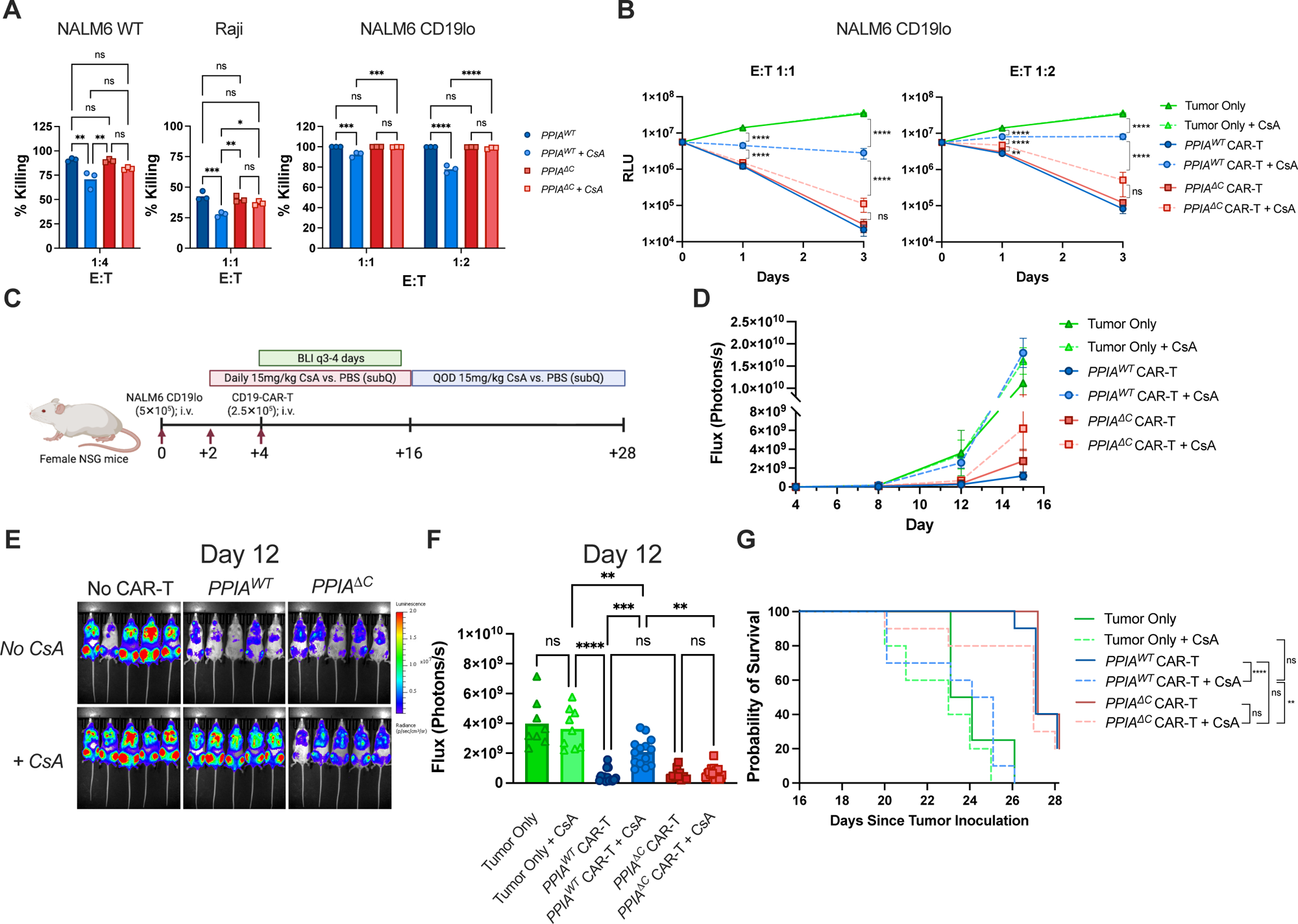
*PPIA*^Δ*C*^ CAR-T cells exhibit enhanced cytotoxicity towards leukemia cells *in vitro* and *in vivo*. A) Three-day cytotoxicity assay for which CD19 CAR-T cells were co-cultured with luciferase positive NALM6 WT, Raji, or NALM6 CD19^lo^ cells at the indicated E:T ratios. B) The NALM6 CD19^lo^ experiment shown in (A) now shown as luminescence over time as a reflection of remaining tumor burden. RLU – Relative Light Units on the luminometer. Group comparisons were made by 2-way ANOVA with multiple comparisons. C) Schematic for *in vivo* assay of CAR-T cell killing of a NALM6 CD19^lo^ leukemia model. D) Absolute Flux over time, where Flux saturates the detector in the 10^10^ range, such that it no longer tracks proportionally with increasing tumor burden. E-F) Representative (E) images of mice in different experimental groups from Day 12 with corresponding (F) Flux analyzed by One-Way ANOVA with multiple comparisons. G) Kaplan-Meier survival curve, with curves compared using a log-rank test with Holm-Sidak correction for multiple comparisons. In vivo studies include 9-15 mice per group, pooling two experiments for Flux, and 5-10 mice per group for survival studies. *<0.05, **<0.01, ***<0.001, ****<0.0001

Next, we evaluated the *in vivo* anti-leukemic efficacy of *PPIA*^Δ*C*^ CAR-T cells in a xenograft model (**Fig. 6C**). NSG mice received daily subcutaneous injections of PBS or CsA, leading to an average trough of 350 ng/mL for the latter. CsA significantly inhibited tumor cell clearance by *PPIA^WT^* CD19 CAR-T cells, as reflected by flux over time (**Fig. 6D**). For example, on Day +12, mice treated with *PPIA^WT^* CAR-T cells had a flux of 2.1 (0.9) x 10^9^ photons/s for those also receiving CsA vs. 5.2 (4.5) x 10^8^ photons/s for those receiving sham injections (p = 0.0001). By contrast, for mice treated with *PPIA*^Δ*C*^ CAR-T cells, flux was comparable in groups receiving CsA vs. sham injections (7.3 (4.0) x 10^8^ vs. 5.7 (3.8) x 10^8^ photons/s, respectively; p = 1.0) (**Fig. 6E-F).** Mice receiving CsA had a significantly increased median survival if they were treated with *PPIA*^Δ*C*^ CAR-T cells vs. *PPIA*^WT^ CAR-T cells (**Fig. 6G**; p = 0.004). Notably, the *PPIA*^Δ*C*^ CAR-T cell + CsA group had similar survival to the CAR-T cell groups receiving sham injections, whereas the *PPIA*^WT^ CAR-T cell + CsA group survival was not significantly different from mice that were untreated (Tumor Only and Tumor Only + CsA groups). These experiments demonstrate a clear benefit of *PPIA*^Δ*C*^ CAR-T cells in preserving *in vitro* and *in vivo* cytotoxic function, culminating in increased survival in a murine leukemia model.

### PPIA^ΔC^ CMV VSTs are resistant to CsA and VCS

We also evaluated the impact of the *PPIA*^Δ*C*^ edit on an additional IEC therapy - VSTs targeting the cytomegalovirus (CMV) pp65 protein.^28^ As shown in **Fig. 7A**, we generated CMV/pp65 specific VSTs that were 13-18% NLV-CMV dextramer positive after a 10-day exposure to pp65 PepMix. After re-stimulation with pp65 PepMix loaded PHA blasts, *PPIA*^Δ*C*^ VSTs demonstrated CD8+ T cell-specific proliferation (**Fig. 7B**) that was better maintained in the presence of CsA and VCS compared to *PPIA^WT^*cells (**Fig. 7C**). For example, for CD8+ VSTs, there was 100 (4)% suppression of division in the *PPIA^WT^*cells vs. 22 (10)% suppression of division with *PPIA*^Δ*C*^ VSTs in the setting of 25 ng/mL CsA (p < 0.0001) and 101 (2)% suppression vs. 9 (15)%, respectively, in the setting of 5 ng/mL VCS (p < 0.0001). Additionally, as observed with *PPIA*^Δ*C*^ CAR-T cells, a higher percentage of *PPIA*^Δ*C*^ VSTs produced IFN-y, TNF-a, and IL-2 when stimulated with PMA/ionomycin or pp65 PepMix with CsA present (**Fig. 7D-E**).

**Figure 7.**
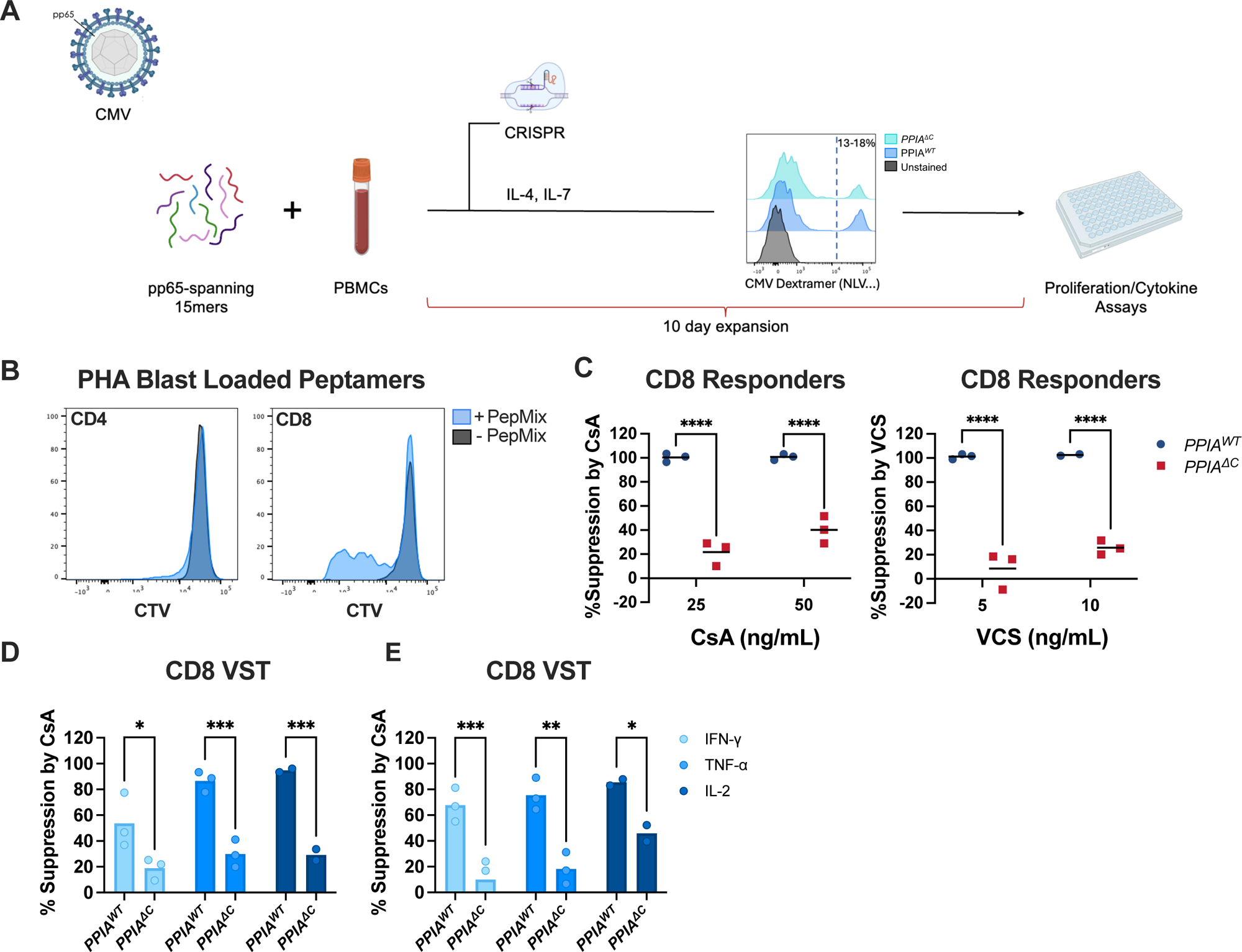
*PPIA*^Δ*C*^ VSTs exhibit preserved division and cytokine production in the presence of drug. A) Schematic for making CRISPR/Cas9 edited CMV specific VSTs from healthy donor PBMCs, of which 13-18% stained positive for HLA-A*02/NLVPMVATV after a first stimulation. B) Representative example of CD8 specific T cell proliferation after restimulation of VST culture with pp65-peptamer loaded PHA-blasts. C) % Suppression of *PPIA^WT^* and *PPIA*^Δ*C*^ VST proliferation in the presence of 25 and 50 ng/ml CsA and 5 and 10 ng/ml VCS. D-E) VSTs were restimulated with (D) PMA/ionomycin or (E) PepMix and stained for intracellular cytokine production. 25 ng/mL CsA was used for these experiments. % Suppression was calculated as described in Methods. A 2-way ANOVA was used for statistical analysis. *<0.05, **<0.01, ***<0.001, ****<0.0001

## Discussion

Here we report a targeted CRISPR/Cas9-based approach to generate CsA-and VCS-resistant IECs by modifying the C-terminus region of CypA protein while preserving *PPIA* expression.

Gene edited CD19 CAR-T cells and VSTs preserved effector and proliferative functions during CsA/VCS therapy, with important clinical implications for IEC use in immunosuppressed patients. Our approach contains the key design feature of targeting the last exon of the *PPIA* gene. This achieves two critical goals. The first is that edits in this region escape NMD, which is generally not triggered when premature stop codons are found in the last exon.^20^ By avoiding NMD, we preserve translation of the majority of the CypA protein, an enzyme with several critical functions in T cells. The second goal was to specifically induce a frameshift in the C-terminus of CypA. This region is important for the interaction of CsA-CypA with calcineurin A while being separate from the catalytic region of CypA, a domain critical for the performance of its homeostatic functions as a peptidyl-prolyl isomerase.^29^

Our approach can be contrasted with prior strategies to engineer CNI-resistant cells. A gene knock-out strategy targeting the tacrolimus-and rapamycin-dependent immunophilin, FKBP12, has been previously published.^11–15,30^ A similar knock-out approach has not been published for *PPIA/*CypA. This may be due to intolerance of a full *PPIA/*CypA knock-out in T cells, given the critical homeostatic functions to which CypA contributes. Indeed, in the Genome Aggregation Database (gnomAD), which contains genetic information from >70,000 individuals, the ‘probability of loss-of-function intolerance’ (pLI) score (ranges from 0 to 1 with 1 being loss-of-function intolerant) is 0.98 for *PPIA*^31^, implying even haploinsufficiency of this protein may be selected against in the human population. Our results with the Exon 4 editing approach were consistent with this, with negative selection of the edit in cultured T cells. Both FKBP12 and CypA are conserved proteins with homeostatic functions. Most studies have described FKBP12’s role in calcium signaling in cardiomyocytes and neurons;^32,33^ with its role in immunity attributed to inhibition of TGF-f3 signaling.^18^ CypA has been demonstrated to have multiple key roles in normal T cell function: it is highly expressed in T cells and functions in protein folding, cell survival, TCR signaling, infection, and inflammation.^9,24,34–39^

An alternative CsA-resistance approach to *PPIA* gene editing has been described, which is to disrupt CsA-CypA docking with calcineurin A by gene addition of a modified calcineurin A (mCNA).^25,40^ However, a limitation observed with this strategy is that cells expressing the mCNA have disrupted downstream NFAT signaling as well as other transcriptomic differences compared to unmodified cells potentially impacting effector function.^25,40^ For this reason, we focused on selective gene modification of *PPIA*/CypA, to avoid both the loss of viability with a total *PPIA* gene deletion and the abnormal NFAT signaling observed with the mCNA addition.

As predicted, we observed preserved cell survival and expansion, with stability of the Exon 5 gene edit for >3 weeks in primary T cells, and no detrimental impact on T cell activation or calcineurin signaling.

As we consider application of the *PPIA*^Δ*C*^ edit to IEC therapies, the differential impact of CsA on 28(versus BB(CAR-T cells deserves discussion. It has been previously demonstrated that strong CD28 co-stimulation may enable T cells to overcome CNI-based suppression, possibly due to decreased NFAT export from the nucleus and enhanced PKC0 signaling.^41–44^

We confirmed this also applies to CAR-T cells, as BB(CAR-T cells were more sensitive to CsA-mediated inhibition than 28(CAR-T cells. However, the latter were still sensitive to CsA when cultured with decreased tumor antigen stimulation and could be rendered resistant to CsA/VCS by the *PPIA*^Δ*C*^ edit. These CsA and VCS resistant CAR-T cells may be considered for post-HCT patients with early B-cell malignancy relapse, HCT or SOT patients with post-transplant lymphoproliferative disorder, or patients with lupus nephritis, who are increasingly treated with VCS.^45,46^

We also highlight the application of the *PPIA*^Δ*C*^ edit to VSTs. Viral reactivation is a leading cause of morbidity and mortality in HCT and SOT patients.^47,48^ In addition to triggering organ dysfunction, there is a bidirectional relationship between viremia and graft-vs-host disease/graft rejection.^49^ Reduction in immune suppression may be used to treat viral reactivation and disease, but further increases the risk of alloreactivity, and traditional antiviral therapies may be insufficient to clear the viral infection and carry significant toxicities.^50^ The demonstration of substantial clinical responses with VSTs not resistant to immunosuppression^51,52^ provides a strong rationale for the utility of VSTs to treat otherwise refractory viral diseases. The addition of CsA/VCS resistance holds the promise of increasing the proliferation, peak expansion, and cytokine production of these cells in HCT and SOT patients treated with CsA or VCS, which may further increase their efficacy.

In conclusion, we have developed a novel strategy for engineering T cell resistance to CsA and VCS. This strategy can be incorporated into diverse cellular products for patients with allo-and auto-immune disease who require ongoing immune suppression with these agents.

## Materials and Methods

### Immune Cell Isolation

Peripheral blood mononuclear cells (PBMCs) were isolated from healthy human donors using a Ficoll gradient. A Pan-T cell isolation kit (Miltenyi) was used to isolate T cells. Cells were cultured in X-VIVO15 (Lonza) media supplemented with 10% fetal bovine serum, 1% penicillin/streptomycin, 2 mM of Glutamax, and 50 µM f3-mercaptoethanol. IL-2 (R&D) was added at 100 IU/mL for initial T-cell stimulation with 1:1 anti-CD3/28 Dynabeads (Gibco) and lowered to 25 IU/mL once Dynabeads were removed. No cytokines were added during functional assays.

### CAR-T Cell Production

Plasmids encoding lentiviral constructs for CD19-CD28-CD3(and CD19-41BB-CD3(CARs were designed through VectorBuilder. Viral packaging was performed as described.^53^ Isolated T cells were stimulated with anti-CD3/CD28 Dynabeads for 2 days prior to transduction.

Transduction was performed via spinoculation using Lentiboost (Mayflower Bioscience).^53^

### Viral-Specific T Cell Production

PBMCs from HLA-A*0201 bearing CMV seropositive donors (n=3) were resuspended with 50 ng/million cells of pp65 PepMix (JPT). After a 60-minute incubation, cells were plated at 1 million cells/mL in a 24-well plate in media containing IL-4 and IL-7 (R&D both at 10 ng/mL).^54^ Fresh media and cytokines were provided every three days. CMV specificity of expanded CD8+ T cells was assessed using the HLA-A*0201-specific dextramer NLVPMVATV (Immudex).

### CRISPR/Cas9 Editing

T cells were CRISPR/Cas9 edited after two days of anti-CD3/CD28 Dynabead stimulation, and CAR-T cells were edited two days after lentiviral transduction. CMV-pp65 peptide stimulated PBMCs were edited three days after exposure to pp65 PepMix. CRISPR/Cas9 editing was performed as described.^55^ Guide RNAs (gRNA) were made by combining tracrRNA (IDT) with negative control cRNA (IDT) or cRNAs for *PPIA* (Exon 4: ATCCTAAAGCATACGGGTCC; Exon 5: GTGTTTGGCAAAGTGAAAGA). DNA was sequenced by Genewiz. Synthego ICE Analysis v3.0 was used to determine editing efficiency.

### DNA and Protein Modelling

Frameshift variants were aligned in TCoffee^56^ and viewed using Mview.^57^ Structural predictions were made using AlphaFold Colab,^22^ with pdb sequences compared to wildtype CypA (PDB:3k0m), viewed in Pymol v.2.5.5.

### CypA Protein Validation

For flow cytometry, cells were fixed with BD Cytofix/Cytoperm and stained with CoraLite®594-conjugated CypA monoclonal antibody (Proteintech; 1:50). For Western blot, membranes were labelled with rabbit CypA polyclonal antibody (1:1000; Fisher; PA1-025) followed by an anti-rabbit HRP secondary antibody (1:5000) using 3% BSA blocking buffer. The membrane was stripped and re-stained with actin polyclonal antibody (PA5-78715). For confocal imaging, cells were fixed with BD Cytofix/Cytoperm, stained with the same polyclonal CypA antibody at a 1:100 dilution followed by an AlexaFluor488 donkey anti-rabbit secondary at 1:250 and imaged on a Zeiss 880 laser scanning confocal microscope.

### RNA-seq

T cells from day 6 after CRISPR editing were either CD8 or CD4 selected using a magnetic column (Miltenyi). Cells were kept unstimulated or re-stimulated with anti-CD3/CD28 beads for 4 hours and processed for bulk RNA-seq. Paired-end next generation sequencing was performed using the Illumina HiSeq platform. Adapter and other Illumina-specific sequences were trimmed using the Trimmomatic package (v0.39).^58^ An alignment index was created using the Kallisto package (v0.50.1)^59^ and the GRCh38 cDNA reference transcriptome. Only “protein coding” and “miRNA” transcripts were retained. Transcripts were normalized to 10,000 counts/sample and transformed with the “numpy.log1p” function.^60^ Clustermaps were created via the Seaborn package (v0.13.2) with the linkage method set to “average” and distance set to “Euclidean”.^61^

### Drug Preparation

CsA and VCS (Millipore-Sigma) powders were reconstituted in dimethyl sulfoxide and diluted for *in vitro* assays. Tacrolimus (Prograf 5 mg/mL) and rapamycin (LC laboratories) were also used for *in vitro* assays. SandImmune^®^ (Novartis) was used for *in vivo* proliferation assays.

### Mixed Lymphocyte Reaction (MLR)

CRISPR/Cas9 edited T cell ‘responders’ were labelled with Cell Trace Violet (CTV; Thermofisher) per manufacturer’s instructions (modification: 1 µL/10 million cells). In a 96-well U-bottom plate, 100,000 edited responder T cells were plated with 300,000 HLA-A*02 disparate, irradiated (30 Gy) ‘stimulator’ PBMCs in a total volume of 200 µL. Cyclosporine, voclosporin, tacrolimus, and rapamycin were added at indicated doses. Assays were read on day 5 after staining for CD4 (L200), CD8 (RPA-T8), and HLA-A*02 (BB7.2). The %suppression by drug was calculated by 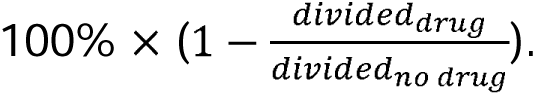^62^

### CAR-T and VST Proliferation Assays

Edited CAR-T cells, 10-12 days after initial stimulation, were stained with CTV and co-cultured with a CD19 positive NALM6 (ATCC) line knocked-out for Beta-2-microglobulin^53^ to focus on CAR-driven proliferation (vs. TCR driven allo-proliferation). Cells were collected for analysis on day 3. Edited VSTs were kept in culture for 10-12 days after initial stimulation, then labeled with CTV, and re-stimulated by co-culture with pp65 PepMix loaded onto autologous PHA blasts.

Samples were collected for analysis on day 4.

### CAR-T and VST Cytokine Stimulation Assay & NFAT inducible proteins

Protein Transport Inhibitor Cocktail (eBioscience) was added to prevent cytokine secretion. For stimulation, either pp65 pepMix was added (for VSTs), NALM6 cells (for CAR-T), or PMA/ionomycin as part of the Cell Stimulation Cocktail (eBioscience; for both VST & CAR-T cells). Intracellular staining was performed after 4 hours using: FITC-anti-human TNF-a (Mab11), BV421-anti-human IL-2 (MQ1-17H12), and BV711-anti-Human IFN-y (B27). 25 ng/mL of CsA was added for VST assays and 100 ng/mL for CAR-T cell assays. The %suppression was calculated as 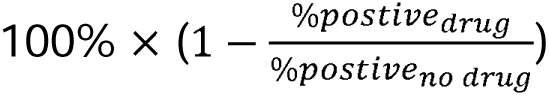 for each cytokine. In separate studies to assess expression of NFAT inducible proteins, CD19-CAR-T cells were co-culture with NALM6 cells at a 4:1 ratio for 2 days and surface stained for CD25 (BC96), CD70 (Ki-24), ICOS (ISA-3), 41BB (4B4-1), OX40 (ACT35), CD40L (24-31), and intracellularly stained for PRF1 (B-D48), GZMB (QA16A02).

### CAR-T Cytotoxicity Assay

Edited CAR-T cells were co-cultured with luciferase positive NALM6 WT, NALM6 CD19^lo^ (kindly obtained from Dr. Robbie Majzner)^27^, or Raji cells (ATCC) at the indicated E:T ratios and durations. %Killing on Day 3 was calculated as 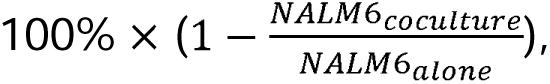 where the numerator and denominator reflect bioluminescent signal. For multi-day experiments, separate plates were created for individual day read-outs, allowing the experiment to continue. Read-outs were performed by adding luciferin reagent from the Bright-Glo Luciferase Assay System (Promega) and measuring luminescence on a EnSpire Multimode Plate Reader (PerkinElmer).

### In vivo studies

Nine-week-old female NOD.Cg-*Prkdc^scid^ Il2rg^tm1Wjl^*/SzJ (NSG) mice were injected with 500,000 NALM6 CD19^lo^ cells on Day 0 by tail vein. On Day +2, mice started receiving either daily subcutaneous injections of 15 mg/kg CsA or PBS (sham). On Day +4, they were injected with 250,000 CAR+ *PPIA^WT^* or *PPIA*^Δ*C*^ CD19-CAR-T cells. Bioluminescence imaging was performed on Days +4, +8, +12, and +15 using D-luciferin (Perkin Elmer) at 150 mg/kg injected subcutaneously and an IVIS imaging instrument. Radiance was calculated using the Living Image software. Mice reached experimental end point when they lost ≥ 20% of body weight, developed a body conditioning score of::: 2, developed clinical symptoms of respiratory distress, or developed neurological symptoms of leukemic infiltration into the central nervous system (gait imbalance, weakness, hind-limb paralysis). A total of n = 9-15 mice were used per group, as indicated, across two experiments for statistical power.

For the *in vivo* proliferation study, healthy NSG mice were injected by tail vein with 5×10^5^ NALM6 WT cells on Day 0 and 6×10^5^ CD19-CAR-T cells on Day +4 after initial labelling with CTV.

As with prior, CsA or PBS was subcutaneously injected daily, starting on Day +2. On Day +8, blood samples were collected via retro-orbital bleed and stained for murine and human CD45 to assess for CAR-T cell division. The %suppression by drug was calculated by 100% x 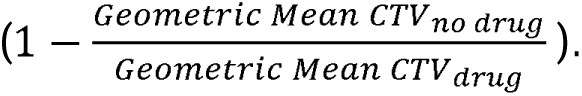

## Data Analysis

Flow cytometry data were analyzed using FlowJo v.10.7.1. Data were graphed using GraphPad Prism v.10, and statistical analysis performed with one-way or two-way ANOVA with multiple comparisons (post-hoc tests specified with each Figure).

## Data Availability Statement

Data can be requested from the authors by reaching out to. The RNA-seq data will be deposited on a public server upon publication.

## Acknowledgments

This work was supported by National Institutes of Health R01-HL095791 (LSK), P01-HL158504 (LSK), U19-AI051731 (LSK), the Lupus Research Alliance (LSK, PAN), P30AR070253 (PAN), R01AR073201 (PAN), R01AR075906 (PAN), R01 HL11879 (BRB), R37 AI34495 (BRB), 2P01 CA065493 (BRB), NIAID T32AI007512-36 (HW), NICHD K12HD052896 Career Development Award (FAC), ASTCT New Investigator Award (VT), Be the Match Foundation/CIBMTR Amy Strelzer Manasevit Research Grant (VT).

## Authorship Contributions

HW, FAC, UG, and LSK designed experiments. HW, JD, KO, XR, EJRP, AA, FW, MW, KAM, GS, WB and LC performed research. HW, RSB, AA, FAC, UG, and LSK analyzed data. HW, UG, and LSK wrote the manuscript. All authors critically reviewed and approved the content of the manuscript.

