## Supplemental Data for "Precision Editing of Cyclophilin A Generates Cyclosporine and Voclosporin Resistant Cellular Therapies"

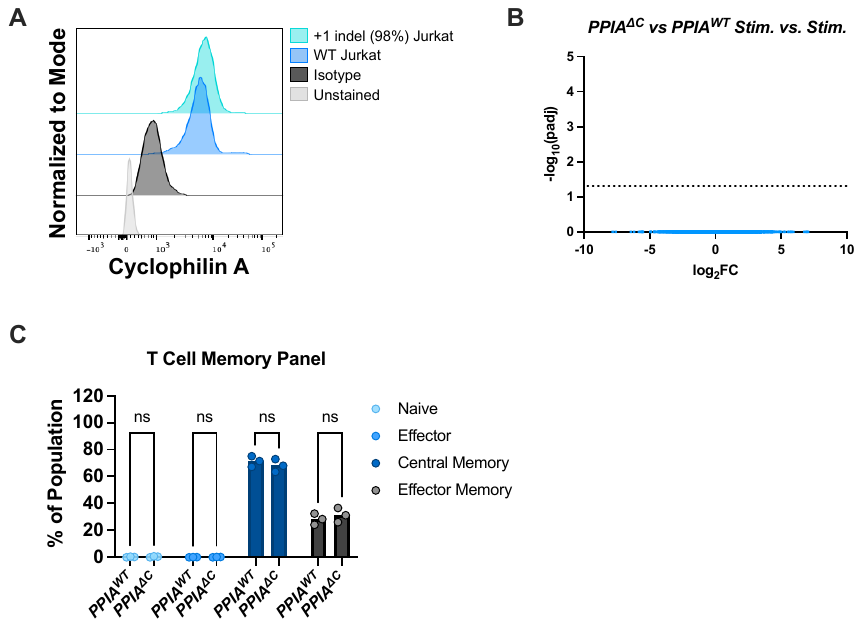

**Supplemental Figure 1.** **Further validation of *PPIA*^Δ^*^C^* cell protein expression and phenotypic characterization.** A) Flow plot demonstrates CypA staining of *PPIA^ΔC^* edited Jurkat cells. Jurkat cells, cultured for 8 weeks at the time of staining, were derived from single cell colonies picked for the +1 bp indel, maintaining a 98% positivity over the culture period. B) Volcano plot of differentially expressed genes in anti-CD3/CD28 bead stimulated *PPIA^WT^* vs *PPIA*^Δ^*^C^* T cells. Based on n = 3 biologic replicates of isolated CD8+ T cells. A dotted line represents the threshold for an adjusted p value of 0.05. C) Memory panel of *PPIA^WT^* vs *PPIA*^Δ^*^C^* T cells after anti-CD3/CD28 bead stimulation, where CCR7 and CD45RO staining was used to define subsets. *<0.05, **<0.01, ***<0.001, ****<0.0001

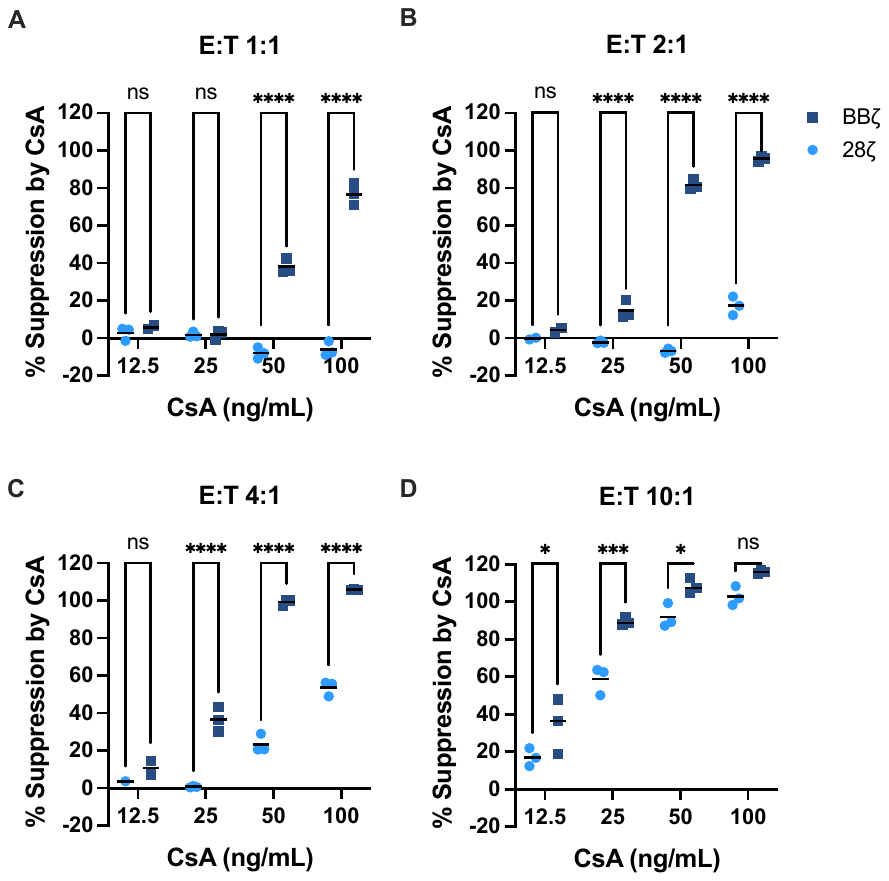

**Supplemental Figure 2.** **BB**$\boldsymbol{\zeta}$ **CAR-T cells are more sensitive to inhibition by CsA than 28**$\boldsymbol{\zeta}$ **CAR-T cells.** Division of CAR-T cells in response to CD19+ $\beta$2M^KO^ NALM6 stimulator cells was assessed based on CTV dilution on Day 3, relative to unstimulated CAR-T cells (negative control). The percent of divided cells was normalized to the drug-free (positive) control after subtracting non-specific proliferation in the CAR-only (no tumor) negative controls for each construct (see Methods$)$. Panels are shown for different E:T ratios of A) 1:1 B) 2:1 C) 4:1 and D) 10:1 with a CsA dose titration. Data shown reflects CAR-T cells gated on CD4. 2-way ANOVA with Sidak post-hoc test used for statistical analysis. *<0.05, **<0.01, ***<0.001, ****<0.0001

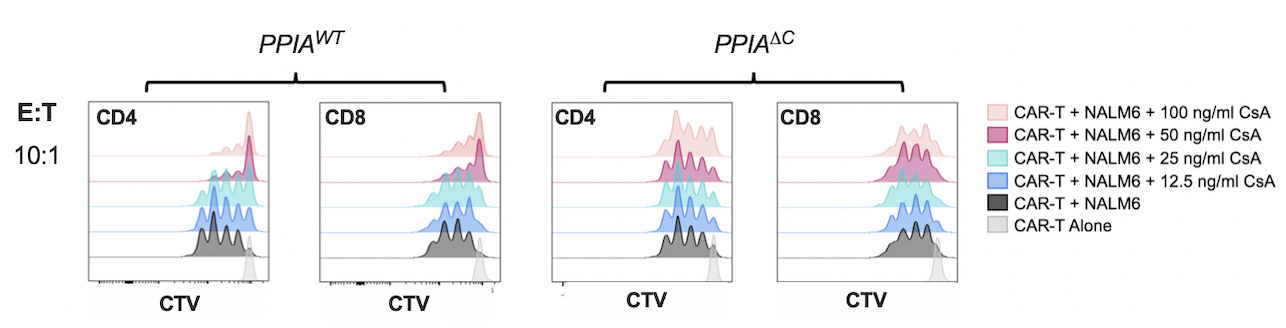

**Supplemental Figure 3: Proliferation of *PPIA^WT^* vs *PPIA*^Δ^*^C^* CAR-T cells in the presence and absence of CsA.** Edited CD19 CAR-T cells were stimulated by NALM6 cells at a 10:1 ratio. The representative division of CAR-T cells is shown on Day 3, indicated by CTV dilution.

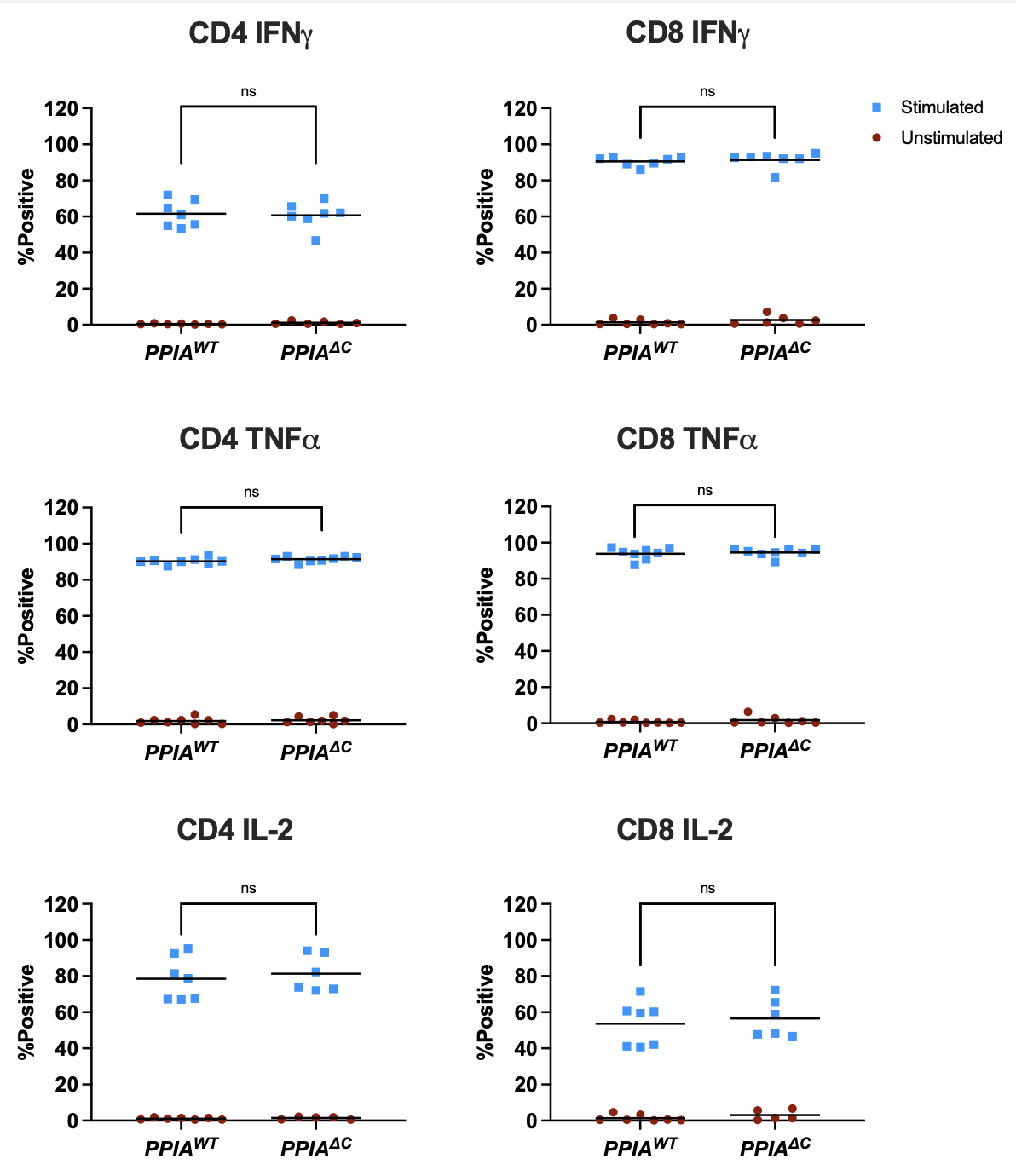

**Supplemental Figure 4**: ***PPIA*^Δ^*^C^* CAR-T cells have equally robust cytokine expression as *PPIA^WT^* CAR-T cells in the absence of drug.** To assess cytokine production, *PPIA^WT^* or *PPIA*^Δ^*^C^* CAR-T cells were stimulated for 4 hours in the presence of protein transport inhibitor +/- PMA/ionomycin and stained for intracellular IFN-$\gamma$, TNF-$\alpha$, and IL-2. 100 ng/mL CsA was used for these experiments. 2-way ANOVA used for statistical analysis.

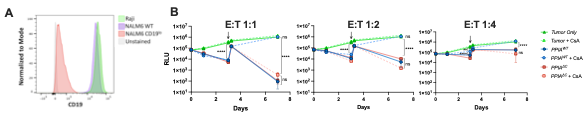

**Supplemental Figure 5**: **Comparison of CD19+ leukemia and lymphoma lines.** A) CD19 expression in different B cell leukemia/lymphoma lines by flow cytometry. B) Long term cytotoxicity assay when CD19 CAR-T cells were co-cultured with luciferase positive NALM6 WT cells, showing bioluminescence over time as a reflection of remaining tumor burden. RLU - Relative Light Units. 40,000 NALM6 cells were reintroduced into the culture system on Day 3 (see arrow) with the corresponding Day 3.25 value imputed based on calculated RLU/cell count. 100 ng/mL CsA was used for these experiments. 2-way ANOVA used for statistical analysis. *<0.05, **<0.01, ***<0.001, ****<0.0001. Experiments based on an n = 3 replicates.
